# Increased oxidative stress tolerance by Superoxide dismutase overexpression in *Mesorhizobium loti*

**DOI:** 10.1101/363556

**Authors:** Pablo J Gonzalez, Mauricio Lozano, Hernán R. Lascano, Antonio Lagares, Mariana N. Melchiorre

## Abstract

Drought and salinity conditions are the major factors affecting nitrogen fixation by legume-rhizobium symbiosis. A global response to these stress conditions is the increase of intracellular ROS leading to activation of antioxidant system to ensure cellular homeostasis. Under the hypothesis that overexpression of *sod* gene in *M. loti* improve tolerance to oxidative stress we decide to investigate the response of constitutive overexpression of the *Mesorhizobium loti* MAFF303099 *sod* gene on tolerance to oxidative stress under free-living conditions. The gene *mlr7636* was overexpressed constitutively under the *nptII* promoter of pFAJ1708 plasmid. Our study revealed that the *sod* overexpressing mutant had five-fold increased SOD activity in periplasmic space and a better tolerance to superoxide and hydrogen peroxide in bacterial killing assays.

**IMPORTANCE:** In this study, we report that the homologous superoxide dismutase overexpression in *Mesorhizobium loti* improves its tolerance to oxidative stress under free-living conditions.

## INTRODUCTION

Reactive oxygen species (ROS) are unavoidable by-products of aerobic life and their signaling role during plant-microbe interactions has been extensively demonstrated. ROS are constantly produced during normal metabolic processes, but their levels are increased under abiotic stress conditions.

In rhizobia, the antioxidant system allows bacteria to modulate ROS levels produced during both the free-living stage and the symbiotic interaction. One of the key enzymes involved in redox modulation of these processes is Superoxide dismutase (SOD, EC 1.15.1.1). SODs are metalloenzymes that act as the primary line of defense against the first free radical produced in the ROS cascade, the superoxide radical (.O_2_^-^), and they have been found in nearly all organisms examined to date (1).

In bacteria, there are three general classes of SODs, which differ in their metal co-factors: the manganese-containing MnSOD (SOD A), the Fe-containing FeSOD (SOD B) and the cambialistic Fe/Mn SODs (CamSOD); the three SODs share high protein sequence and structure similarity (2, 3).

Rhizobiales have shown variability in their superoxide dismutase enzymes: in *Sinorhizobium meliloti* Rm5000, *sodA* is the only gene encoding an active cambialistic SOD with either manganese or iron as co-factor (2). In *Bradyrhizobium japonicum* USDA 110, four genes encoding superoxide dismutases have been reported (genome.microbedb.jp/rhizobase). Two of these genes, *bll7559*, a Fe-Mn SOD (ChrC), and *bll7774* SOD, *sodF*, have been reported as inducible under drought stress (4).

In *Mesorhizobium loti* MAFF 303099, SOD is encoded by the *mlr7636* gene and has been described as a Mn/Fe co-factor binding protein (genome.microbedb.jp/rhizobase). SOD activity (and other cellular processes) produces hydrogen peroxide. In *M. loti*, hydrogen peroxide scavenging occurs through the action of two different catalases, a monofunctional catalase and a bifunctional catalase-peroxidase encoded by *mlr2101* and *mlr6940*, respectively (*genome.microbedb.jp/rhizobase*).

It has been reported that disruption of the *sod* gene in S. *meliloti* affects its symbiotic properties with alfalfa, with the mutant bacteria being affected in the initiation of nodulation, infection and bacteroid development (1). However, to our knowledge, there are no reports about the effect of bacterial SOD overexpression on tolerance to oxidative and salt stress conditions. In this work, a SOD overexpressing *M. loti* was obtained, and its free-living phenotype was investigated. Our results suggest that *sod* overexpression in *Mesorhizobium loti* improves its survival during oxidative stress.

## RESULTS

### Effect of saline and osmotic stress on *M. loti* growth

The effect of saline and osmotic stress conditions on *M. loti* overexpressing *sod* gen (sod1708), the wild-type strain (WT) and the transformed strain with the empty vector (1708) were evaluated in YEM media (control), YEM containing 150 mM NaCl and 15% PEG. Under control conditions, the three strains had equivalent growth rates, whereas in saline treatment, all strains showed slightly higher growth rate than in control condition. PEG treatment showed a 50% decrease in bacterial growth rate, demonstrating that the strains were sensitive to osmotic stress (Fig. 1).

**Figure 1.**
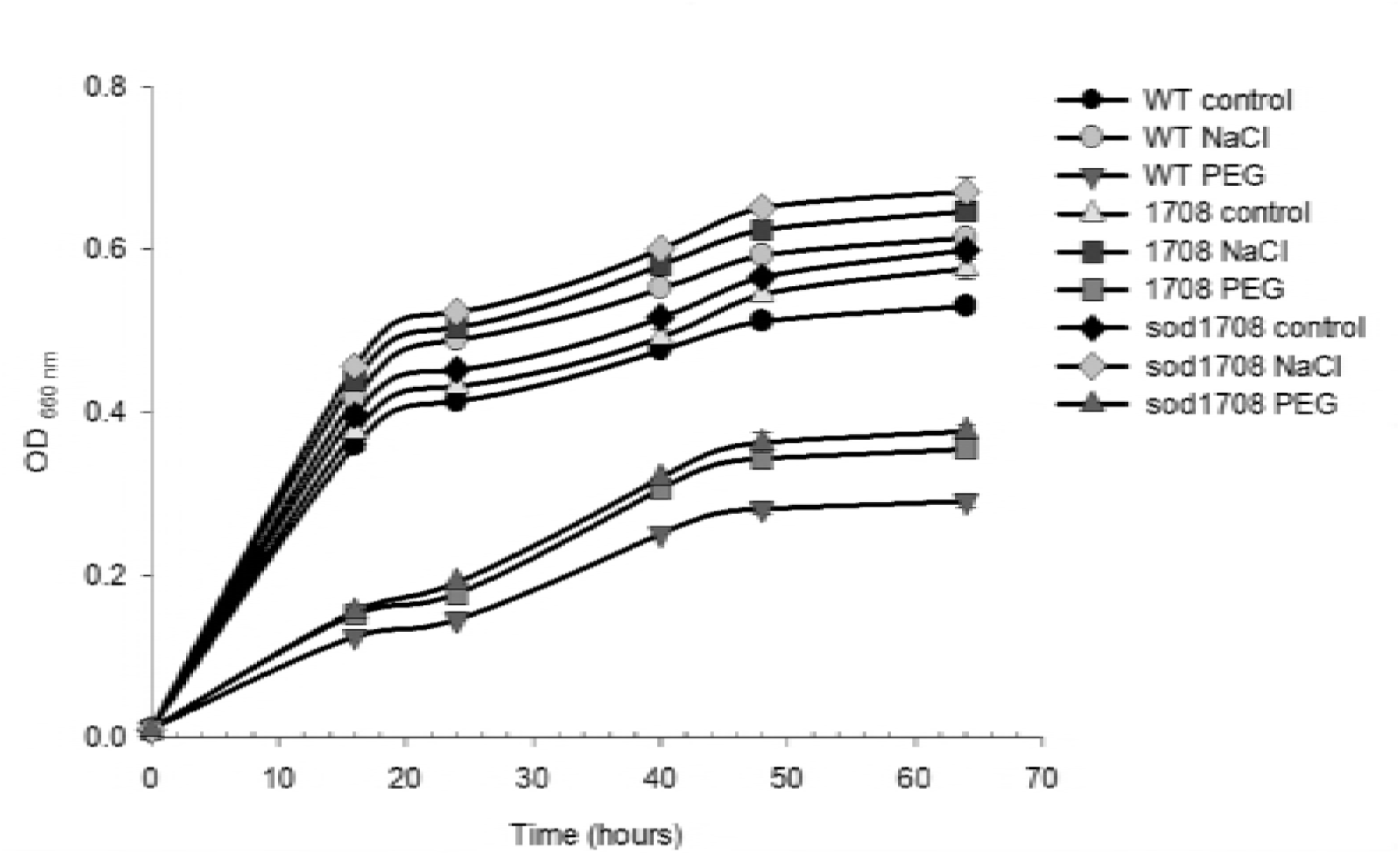
Growth curves of the WT, 1708 and sod1708 strains under osmotic and saline stress conditions. YEM medium was supplemented with 15% PEG or 150 mM NaCl, both of which lowered water potential to −0.84 Mpa. Data are the mean of three independent replicates with the error bars indicating the standard error.

### Effect of saline and osmotic stress on SOD and CAT activities

To evaluate the effect of salt and osmotic stress on the bacterial SOD and CAT, total activities were determined in whole extracts from cells grown under 150 mM NaCl and PEG 15%. Under salt stress, the WT strain showed higher levels of SOD and CAT activities than in control conditions, with approximately 4.8 and 2.8-fold increments, respectively (Table 2). However, under osmotic stress condition, only CAT showed increased activity, whereas SOD decreased. The overexpression of *mlr7636* gene in sod1708 strain led to 3.4-fold increase in SOD activity with respect to WT strain and 3.6-fold with respect to 1708 strain under control conditions. Under salt stress condition, no differences in SOD activity were found between the overexpressor sod1708, WT and 1708 strains (Table 2). Similar responses were found under PEG-induced osmotic stress treatment, with a 4.1- and 4.5-fold increase in SOD activity being detected in sod1708 strain compared with WT and 1708 strains, respectively.

**Table 1.**
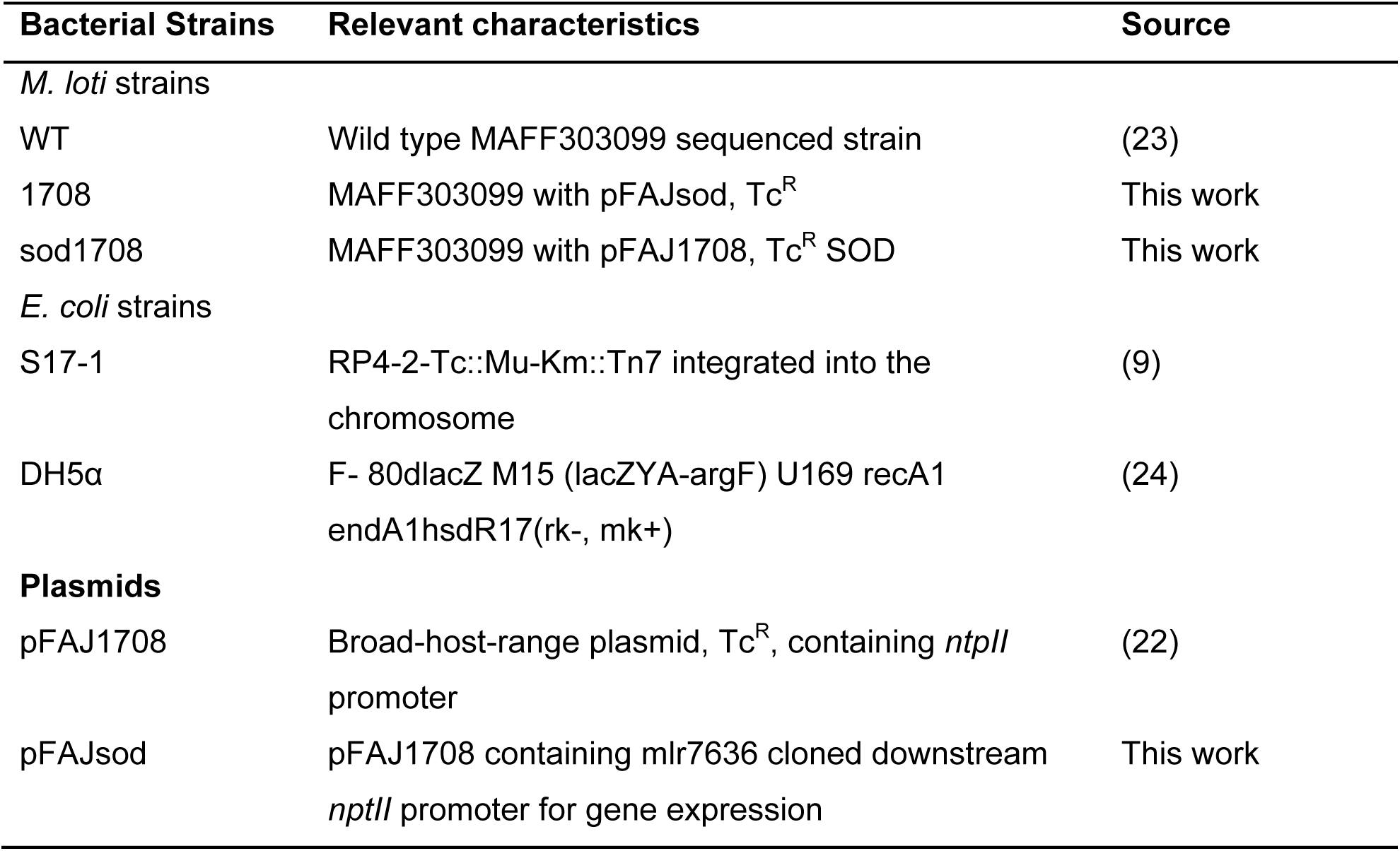
Bacterial strains and plasmids used in this study.

**Table 2.** Specific activities of SOD and CAT of WT, 1708 and sod1708 strains, under NaCI and PEG conditions.

| Strains | SOD activity |  |  | CAT activity |  |  |
| --- | --- | --- | --- | --- | --- | --- |
|  | Control | NaCl | PEG | Control | NaCl | PEG |
| WT | 0.20±0.04a | 0.97±0.11a | 0.10±0.01a | 0.13±0.05a | 0.37±0.09a | 0.58±0.08a |
| 1708 | 0.19±0.04a | 1.01±0.09a | 0.09±0.03a | 0.16±0.04a | 0.36±0.02a | 0.54±0.06a |
| sod1708 | 0.69±0.06b | 1.16±0.13a | 0.41±0.05b | 0.45±0.11b | 0.68±0.10b | 0.74±0.05a |

Results of enzyme activities are presented in U (μg protein)<sup>-1</sup>
Each value is expressed as mean ± SE (n=3)
Different letters in each column indicate significant differences ( $p \leq 0.05$ DGC test)

CAT activity increased significantly in sod1708 strain in control conditions and saline stress treatment, but it showed no differences from the osmotic treatment.

### Increase of periplasmic SOD activity in *M. loti* sod strain

Specific SOD activity determined at an early growth stage (DO_600_ ~0.1) was lower than at the late exponential phase; however, no differences in whole cell lysates were found between strains. The analysis of specific SOD activity in the periplasmic fraction revealed that sod1708 strain showed a 5-fold increase of enzyme specific activity compared to non-overexpressing strains (Fig. 2). In order to assess the cytosolic contamination of the periplasmic extract, the specific G6PDH activities of both the whole cell extract and the periplasmic fraction were determined (Fig. 3). The results showed low specific G6PDH activity levels for the periplasmic fractions with less than 12, 6 and 2% of the specific activity of the whole cell lysate for WT, 1708 and sod1708 respectively.

**Figure 2.**
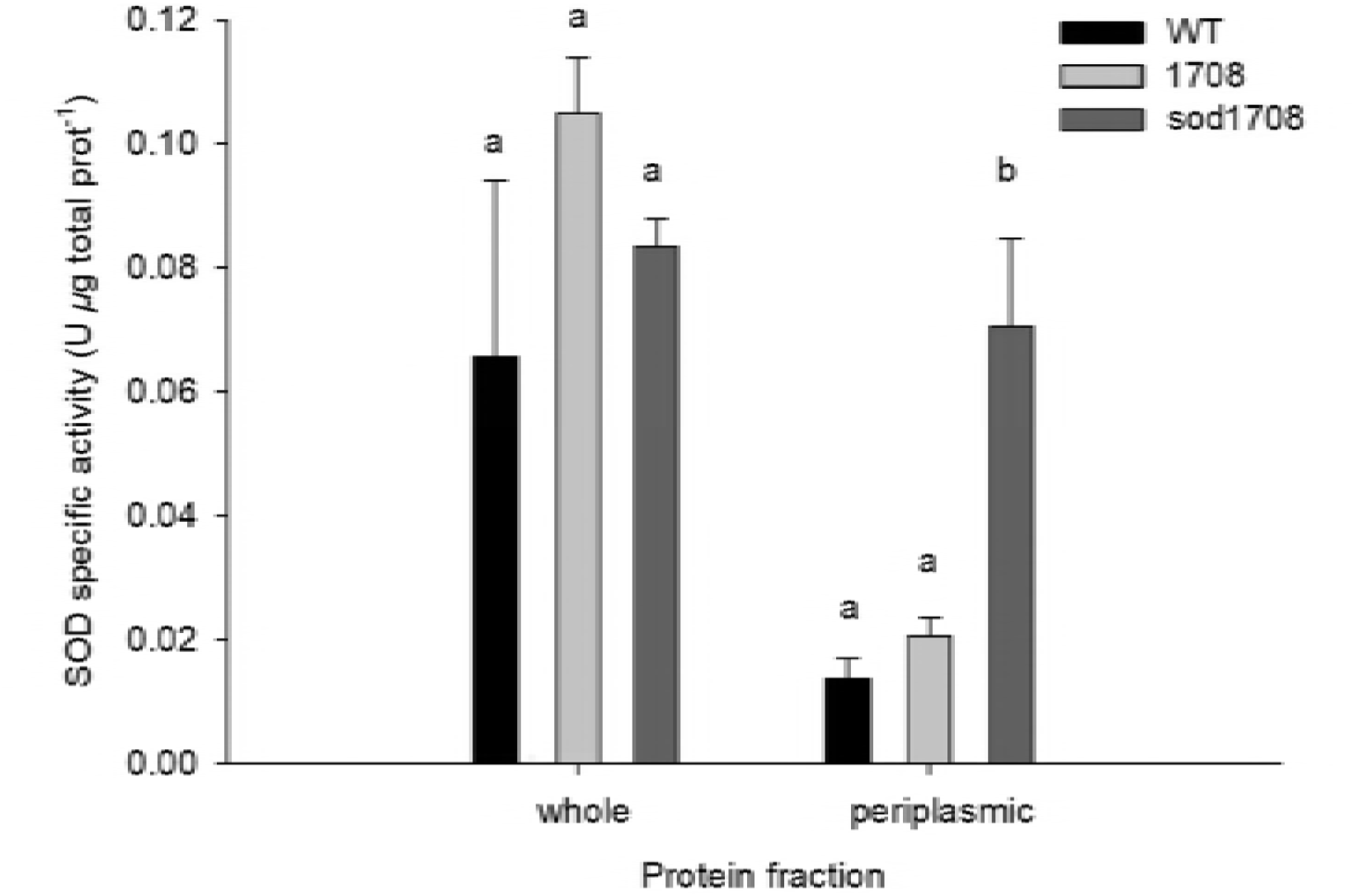
Specific SOD activity of the WT, 1708 and sod1708 strains in whole and periplasmic extracts. Data are representative of at least three independent replicates. Different letters indicate significant differences in the mean (*p*≤0.05 DGC test).

**Figure 3.**
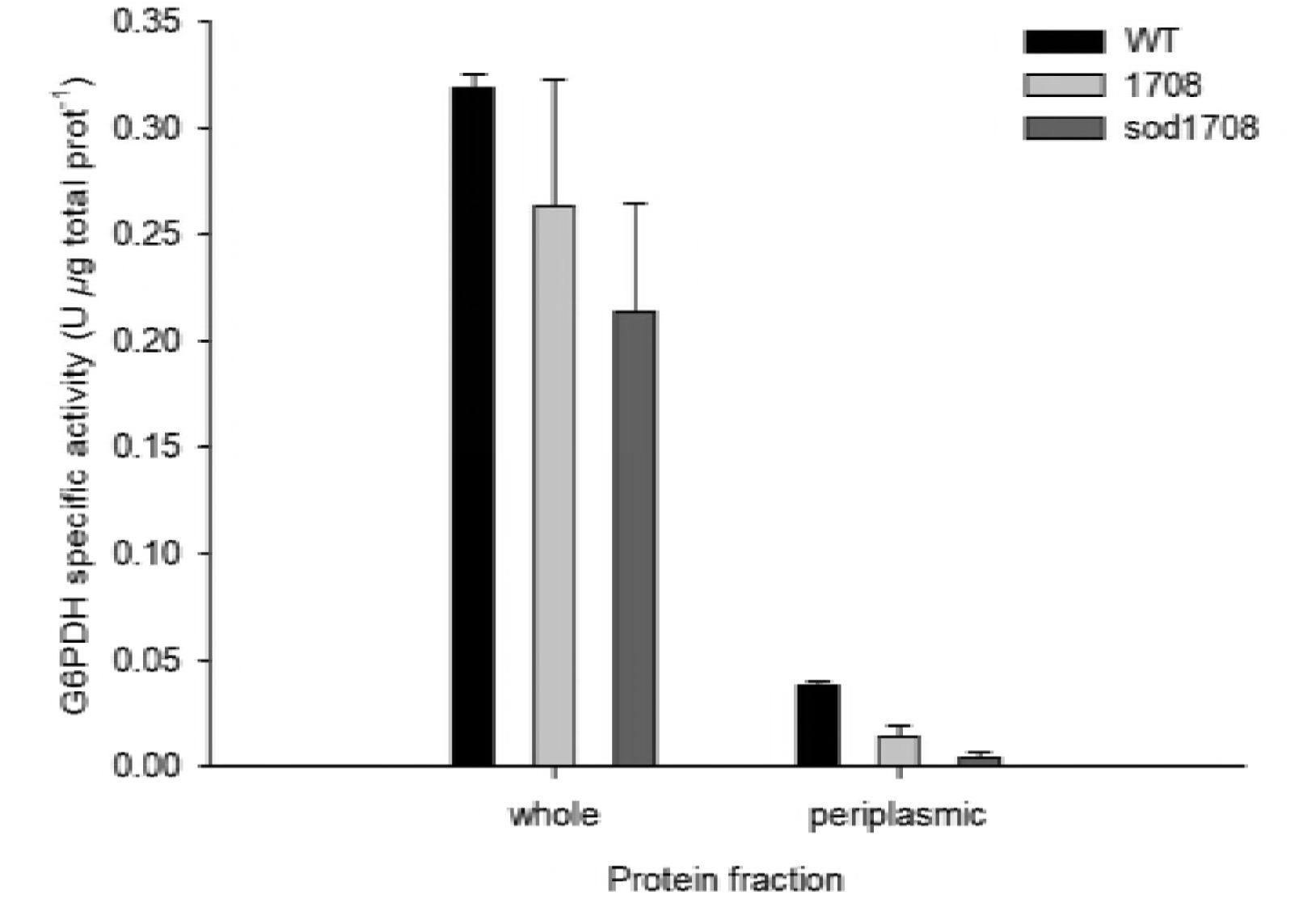
G6PDH activity as cytoplasmic marker and control of contamination in periplasmic preparation of the *M. loti* strains. Data are representative of three independent replicates. Error bars represent the standard error.

### Isoforms and localization of SOD in *M. loti* sod

The expression of SOD isoforms revealed in zymograms using whole cell and periplasmic protein fractions showed changes in activity patterns among the WT, 1708 and sod1708 strains. A unique activity band, which we named basal sod (BSOD) in this report, was observed in whole cell extracts of all strains, whereas the sod1708 strain showed three extra bands (Fig. 4 A). In periplasmic fractions, BSOD was absent in WT, whereas the sod1708 and the 1708 strains shared the same activity profile with similar relative mobility (Rf) to those found in whole cell extracts, including the three extra bands. Moreover, the three extra activity bands were inhibited by 10 mM H_2_O_2_, indicating that these isoforms used iron as cofactor (Fig. 4 B).

**Figure 4.**
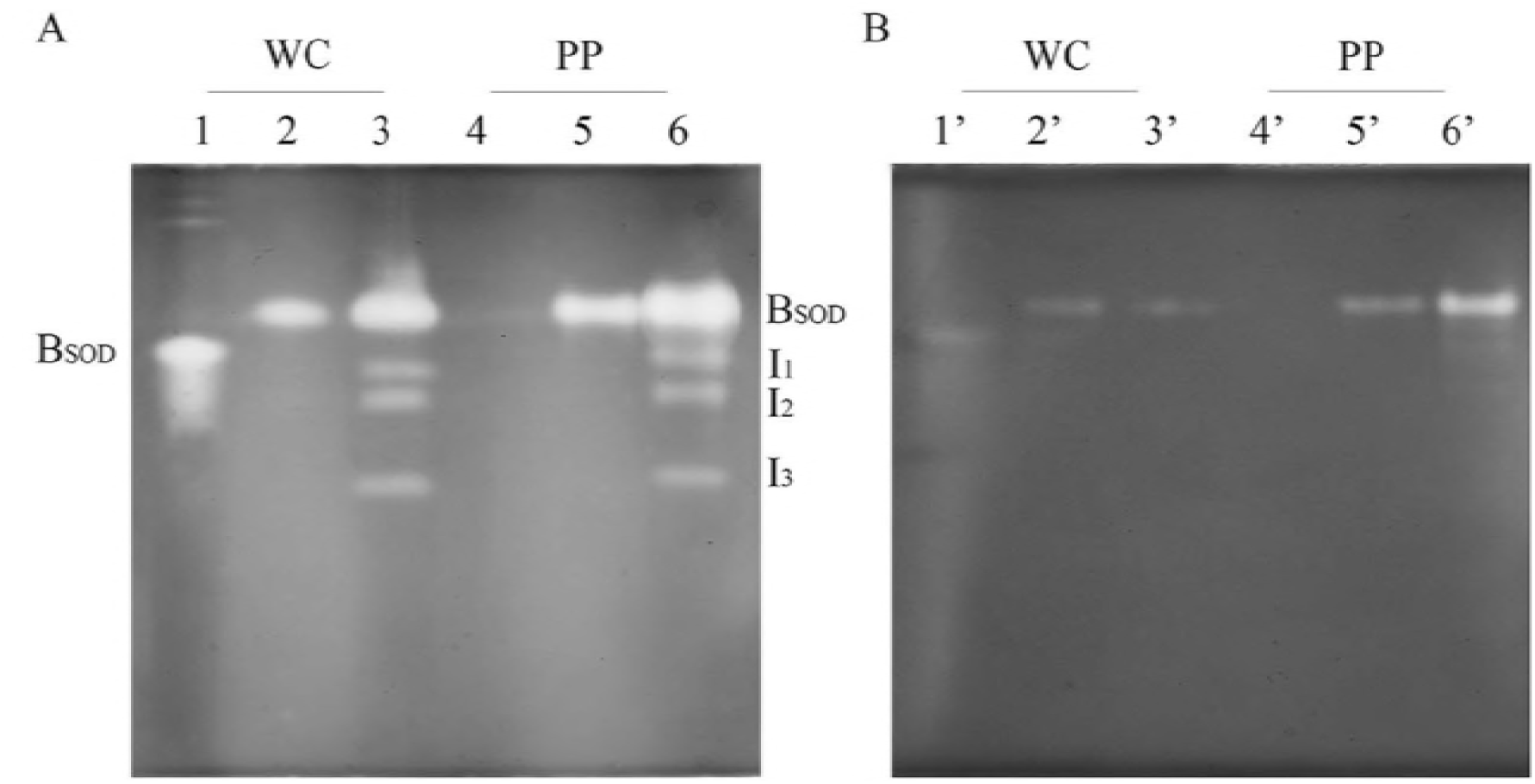
A- Total SOD activity revealed by ND PAGE of the WT, 1708 and sod1708 strains. Lanes 1-3 correspond to whole cell (WC) and lanes 4-6 correspond to periplasmic (PP) extracts. B- SOD Inhibition by 10 mM H_2_O_2_.

**Figure 5.**
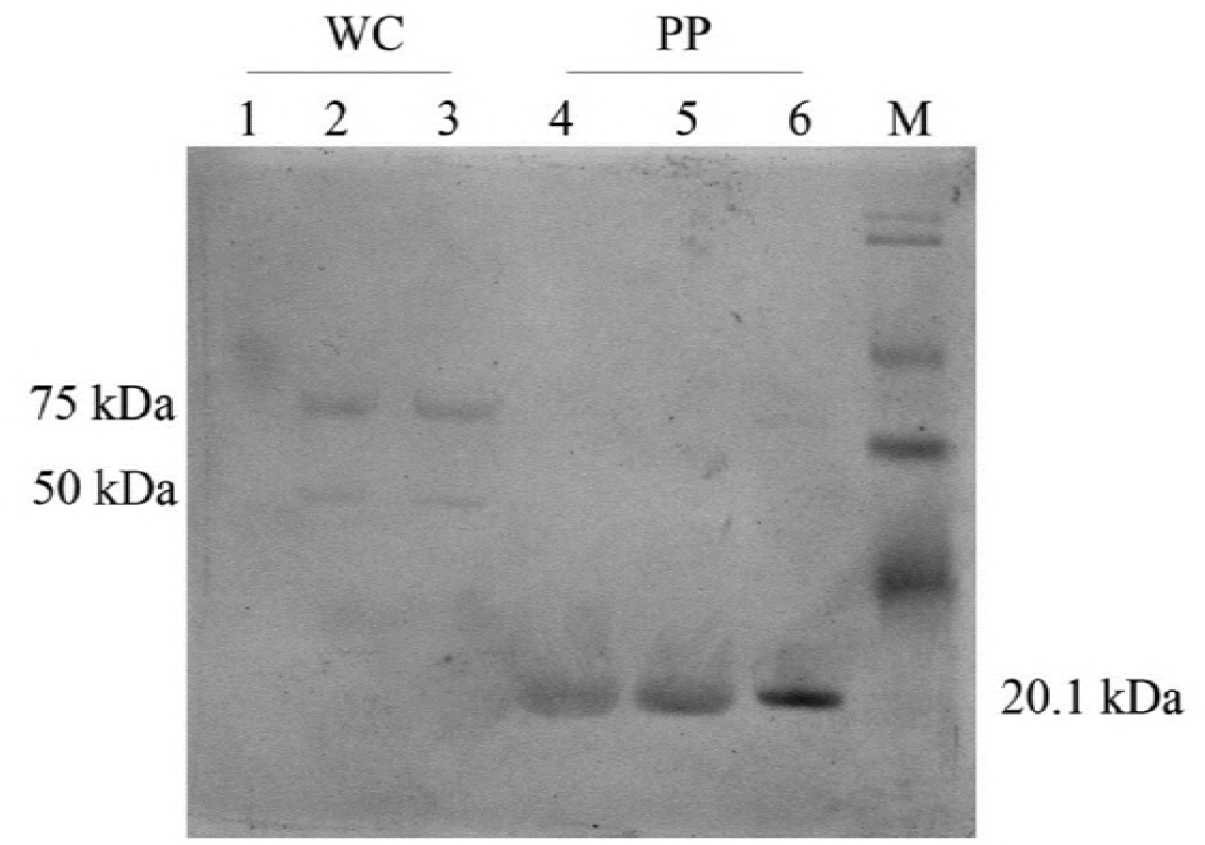
Western-blot analysis using anti-FeSOD antibody of the WT, 1708 and sod1708 strains. Lanes 1-3 correspond to whole cell (WC), lanes 4-6 correspond to periplasmic (PP) extracts. Lane M corresponds to prestained SDS-PAGE broad range standard (Bio-Rad).

### WESTERN-blot analysis

Western blot analysis with FeSOD antiserum detected bands of 20.1 kDa in periplasmic extracts from all strains. On the other hand, this band was not detected in whole cell extracts. However, the 20.1 kDa protein in periplasmic extracts is not in agreement with the theoretical molecular weight of 22.7 kDa predicted for the protein encoded by *mlr7636* gene. Therefore, the difference in 2.6 kDa mass obtained in the periplasmic extracts could be due to scission of a signal peptide. Nevertheless, the presence of signal peptide for Sec or TAT pathway could not be predicted in the first 70 amino acids of the N-terminal end of the amino acid sequence.

### Bacterial tolerance to superoxide and H_2_O_2_ in killing assay

To assess the effect of the SOD overexpression on *M. loti* under excess of reactive oxygen species production, we performed a bacterial killing assay using cultures of sod1708, WT and 1708 strains, subjected to exogenously generated .O_2_^-^ by the addition of hypoxanthine and xanthine oxidase. The results showed a 20% reduction in CFU/mL in sod1708 strain after 30 min of treatment. This survival reduction was not statistically significant with respect to the untreated control and no further decrease was recorded up to 120 min (Fig. 6 A). In contrast, both WT and 1708 strains showed more than 90% reduction in viability compared to the untreated control 30 min after .O_2_^-^ treatment, with no viable cells being detected even after 60 min. Catalase supplementation (1 U/mL) to the hypoxanthine/xanthine oxidase system restored the growth of the overexpressing sod1708 strain to similar levels to those of the untreated control (Fig. 6 B). On the other hand, exogenously added catalase restored the CFU/mL values to those of untreated cells up to 30 min, whereas a decrease of approximately 55% was observed starting at 60 min.

**Figure 6.**
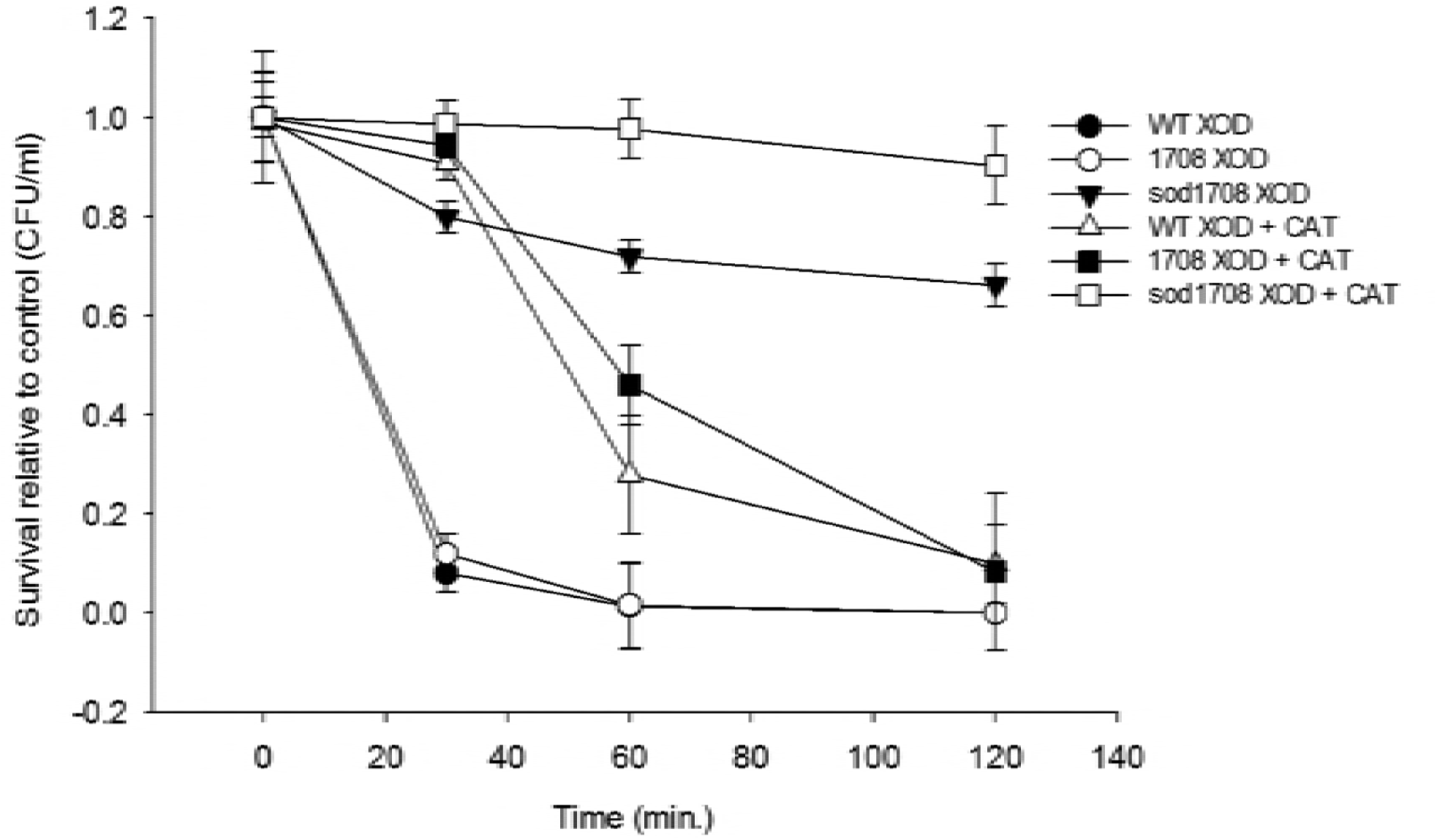
A- Survival of the sod1708 strain to superoxide anion in killing assay (.O_2_^-^ generating conditions). B- Survival in killing assay with addition of 1U/mL catalase. Data are the mean of three independent replicates. Error bars represent the standard error.

To test whether increased superoxide tolerance of the sod1708 strain may be due to enhanced H_2_O_2_ tolerance, survival percentage was determined by adding 1 mM H_2_O_2_ to the growth media. Survival of *M. loti* sod strain was not significantly affected compared with the untreated control up to 60 min, whereas at 120 min only 26% of bacteria survived. In contrast, the CFU/mL of the WT and 1708 strains showed a reduction of 27% and 90% after 30 and 60 min, respectively, of hydrogen peroxide treatment (Fig. 7).

**Figure 7.**
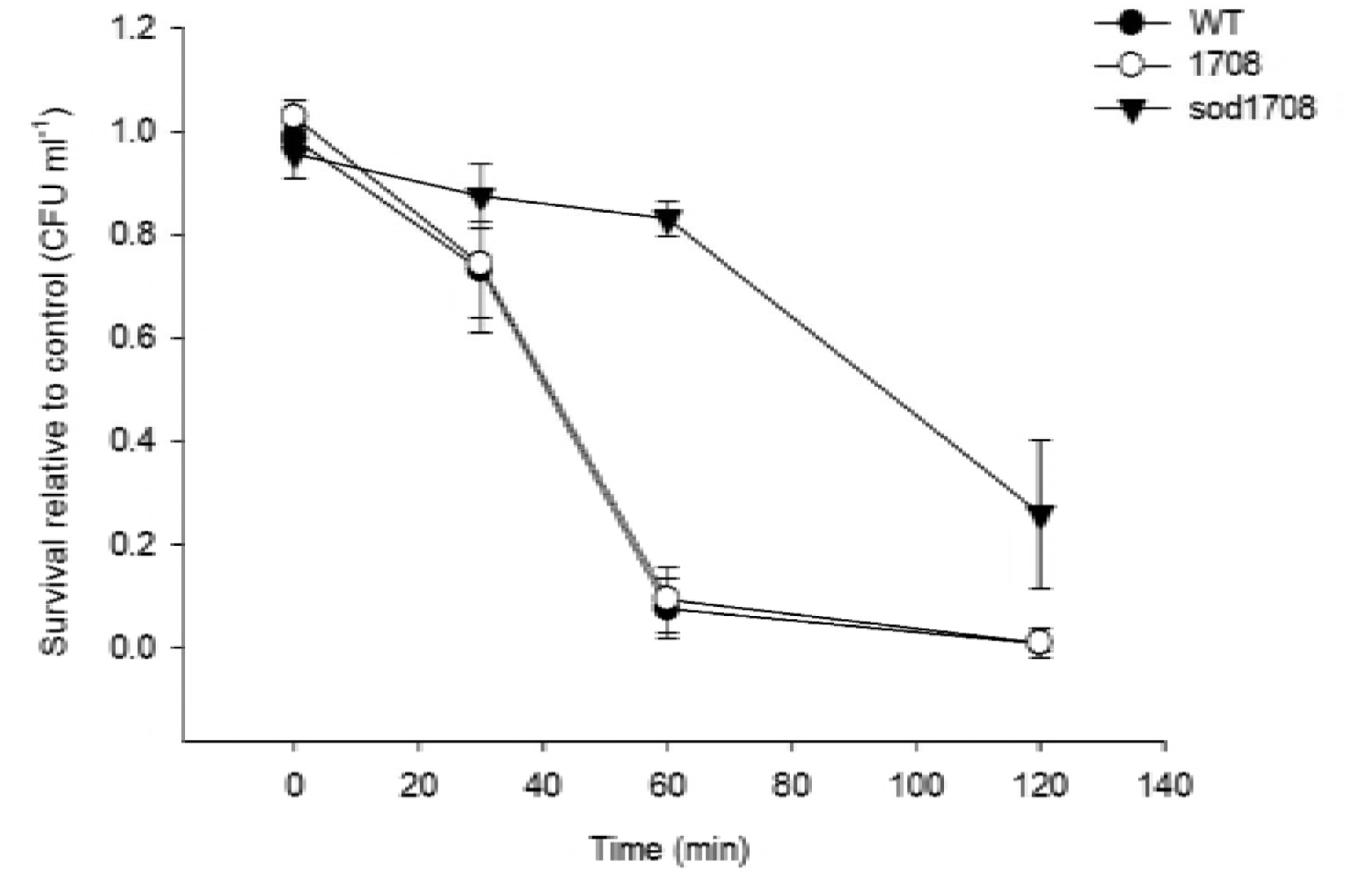
Survival of the sod strain to H_2_O_2_ 1 mM. Data are the mean of three independent replicates. Error bars represent the standard error.

### Phylogenetic analysis of rhizobia *Sod* genes

The consensus phylogenetic tree with 11 SOD sequences from Rhizobase database, that had ≥50% identity with Mlr7636 sequence, showed two clusters (Fig. 8). Cluster I includes the product of the *mlr7636* gene between putative *SodB* of *Rhizobium leguminosarum* bv. viciae 3841 and *SodB* of *Rhizobium etli* CFN42 (RHE CH01203). Moreover, this subgroup was related to *SodB* FeSOD of *Sinorhizobium meliloti* 1021 (SMc00043) and *SodB* Mn-SOD of *Rhizobium sp*. NGR234 (NGR c07300). Also in cluster I three SODs capable to use Fe or Mn as co-factor were located: SODB from *Bradyrhizobium sp*. BTAi1 (BBta 1335), SOD of *Bradyrhizobium sp*. ORS278 (BRADO6273) and SOD of *B. japonicum* USDA110 (bll7774). Cluster II includes sequences that showed lower identity levels with the Mlr7636 sequence, and it was composed by FeSOD of *K. pneumoniae* Kp342 (GKPORF B1093), SOD of *C. taiwanensis* LMG19424 (RALTA A0566) and Fe/Zn/MnSOD of *Azospirillum sp*. B510 (AZL014560).

**Figure 8.**
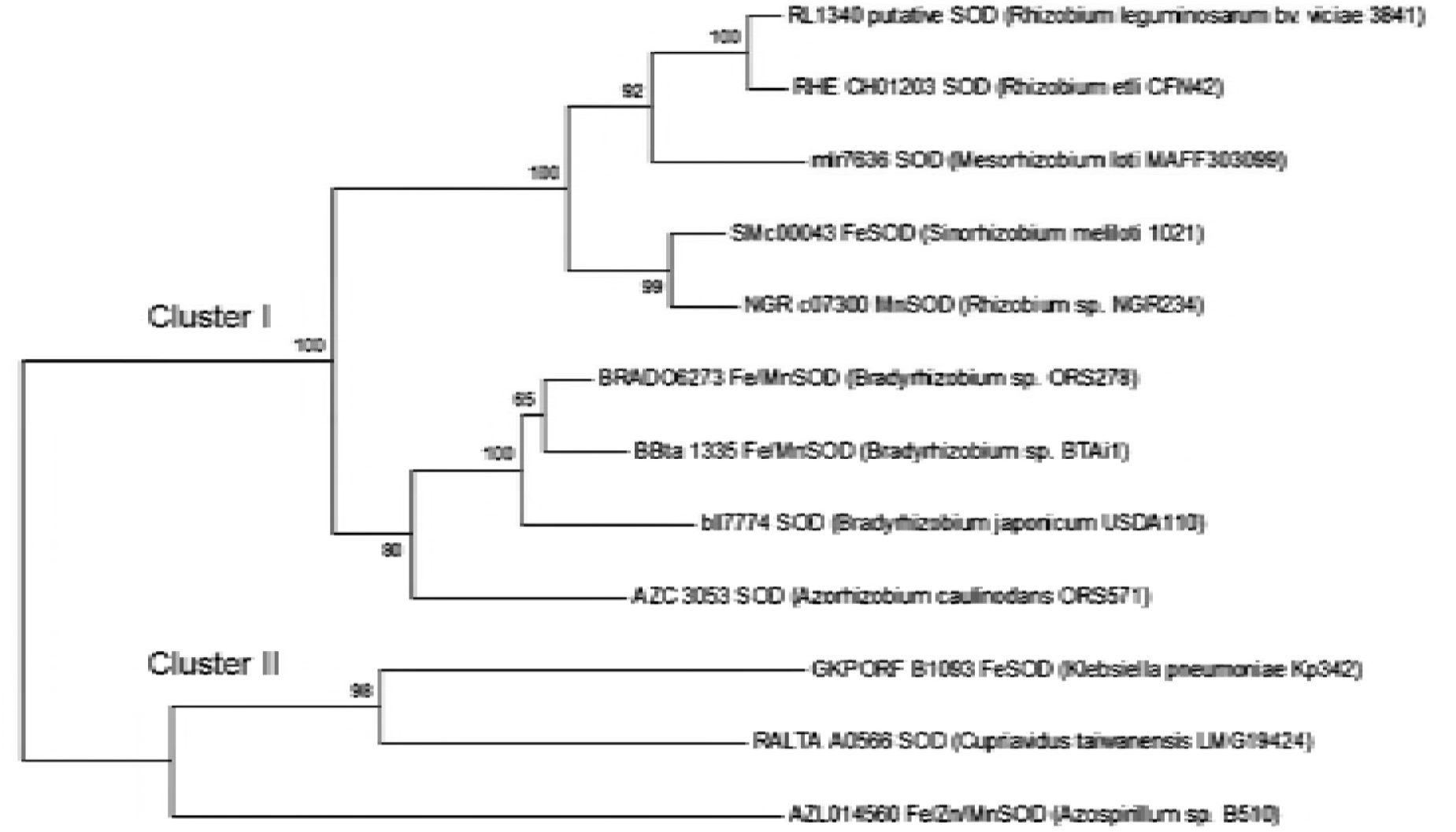
Neighbor-joining phylogenetic tree based on aligned sequences of 11 sod amino acid sequences from the Rhizobase database sharing more than 50% identity with the *mlr7636* product.

## DISCUSSION

The genome of *M. loti* MAFF303099 includes the *mlr7636*, *mlr2101* and *mlr6940* genes, which encode a superoxide dismutase, a catalase and a catalase / peroxidase, respectively (genome.microbedb.jp/rhizobase). The complex detoxification mechanism of ROS in *M. loti* MAFF303099 involves the catabolism of superoxide and H_2_O_2_, therefore, superoxide dismutase and catalase are two of the main enzymes involved in superoxide and hydrogen peroxide catabolism.

Our results showed that exposure of *M. loti* to saline stress increased both SOD and CAT activities. Moreover, under osmotic stress, SOD activity was reduced and CAT presented the highest induction in response to both saline and osmotic stress. Induction of SOD and CAT activities under stress conditions are in agreement with results reported by Cytryn *et al*. (4), who described the induction of genes of *B. japonicum* exposed to reduced water activity.

### Overexpression

The sod1708 strain showed 3.4 fold increased SOD activity in comparison with the WT and 1708 strains under control conditions, whereas no differences were found between the strains when they were exposed to osmotic and salt stress. SOD was detected in the periplasmic fractions from the three strains, but sod1708 strain showed higher SOD activity than the non-overexpressing strains, indicating that it was exported to the periplasmic space. Furthermore, zymograms revealed the presence of three additional SODs in the overexpressing *M. loti* sod strain. These bands were inhibited by H_2_O_2_ indicating that they were FeSOD (21). In contrast, lack of inhibition with H_2_O_2_ demonstrated that the common BSOD corresponded to a MnSOD. It is difficult to explain why the BSOD migration of 1708 and sod1708 were somewhat different from the wild type strain, although it could be due to metabolic changes occurring in transformed cells (22).

The homologous gene overexpression as well as the multiple activity bands differentially inhibited by H_2_O_2_ in zymograms, support the idea that SOD from *M. loti* MAFF303099 could have a cambialistic behavior able to form a complex with either Fe or Mn (2), further investigation will be required to elucidate this aspect.

On the other hand, the Western blot analysis revealed the presence of single bands of approximately 20.1 kDa in periplasmic extracts of all strains. In addition, the calculated molecular weight of the detected protein was 2.6 kDa lower than the theoretically predicted for the *mlr7636* product. Although the signal peptide prediction showed no detectable Sec or TAT signal peptide (18, 19), the difference could be due to scission of a signal peptide during translocation to periplasmic space using an alternative mechanism as it was described in *R. leguminosarum* bv. viciae (10). The consensus phylogenetic tree of Mlr7636 revealed that it was highly related to *R. legumnosarun* bv. viciae 3841, and clustered with FeSOD of *Sinorhizobium meliloti* 1021 (SMc00043) as well as with MnSOD of *Rhizobium* sp. NGR234 (NGR c07300) sequences. The other sub group in cluster I included sodB from *Bradyrhizobium sp*. BTAi1 and BRADO6273 from *Bradyrhizobium sp*. ORS 278; both of them can use Fe and Mn as co-factor.

### SOD overexpression protects against oxidative stress by inducing catalase activity

The protective effect of superoxide dismutase overexpression against oxidative stress in free-living *Mesorhizobium loti* has been demonstrated in killing assays. In the presence of exogenously generated superoxide, the sod1708 strain showed substantially higher tolerance than the controls WT and 1708 strains. The CFU reduction of the *M. loti* sod strain was 20% under oxidative stress condition caused by superoxide, whereas the control strains were non-viable.

The dismutation of superoxide anion by SOD produces H_2_O_2_ and water, so that the addition of catalase to the hypoxanthine/xanthine oxidase reaction in the bacterial killing assay restored the initial CFU values to those reached by the untreated strain. A similar effect was observed in control strains when catalase was added, but in these cases the restored CFU values reached only 45%. These results clearly show that the resulting H_2_O_2_, from superoxide dismutation induced by SOD activity is more lethal on bacteria than superoxide anion per se. Furthermore, our data suggest that homologous superoxide dismutase overexpression increases tolerance to oxidative stress in *M. loti* because of the high SOD activity *per se* contributes to fast superoxide dismutation to H_2_O_2_ and O_2_, and additionally the induced catalase activity reduces the hydrogen peroxide levels.

## MATERIALS AND METHODS

### Bacterial strains and growth conditions

*Mesorhizobium loti* MAFF303099 (WT), 1708 and sod1708 strains were grown in YEM medium (5) aerobically at 28°C. For saline and osmotic stress conditions, YEM medium was supplemented with 150 mM NaCl and 15% (W/V) polyethylene glycol 8000 (PEG) to obtain a water potential equal to −0.84 MPa. Values of water potential were measured using isopiestic thermocouple psychrometry (Wescar^®^ Point Microvoltmeter HR-337 Dew). *Escherichia coli* DH5α and S17-1 strains and were grown Luria-Bertani (LB) (6). Strains harbouring plasmids pFAJ1708 and pFAJsod were grown on Tetracycline 10 μg/mL supplemented medium. The bacterial strains and plasmids used are listed in Table 1.

### Growth kinetics

Cell growth was evaluated in triplicate using a pre-inoculum of 0.8 (OD_600_) with an initial OD_600_~ 0.01 in 50 mL fresh media at 28°C in an orbital shaker (160 rpm). Growth was monitored at OD_600_ in a DU-640 Beckman spectrophotometer (Beckman Coulter, USA) for 64 hours.

### Determination of tolerance to oxidative stress

Tolerance to .O_2_^-^ was determined by conventional bacterial killing assays, in which bacteria were exposed to exogenous .O_2_^-^ generated by the oxidation reaction of xanthine by Xhantine oxidase enzyme (7). Cells were grown up to 0.5 OD_600_ in YEM; then they were centrifuged and resuspended at 10^9^ CFU/mL in sterile phosphate-buffered saline (PBS) and treated with 250 μM hypoxanthine and 0.1 U/mL of Xanthine oxidase. In order to discriminate the effect of .O_2_^-^ and the H_2_O_2_, produced by SOD activity, bovine catalase (1 U/mL) was added to hypoxanthine/xanthine oxidase reaction to reduce the H_2_O_2_ presence and subsequent Fenton chemistry, and to ensure that killing of bacteria was only in response to .O_2_^-^ radicals. The number of viable bacteria was determined at 0, 30, 60, and 120 min of incubation by plating serial dilutions on YEM agar plates. A similar protocol was used for examining the susceptibility to H_2_O_2_. Cultures were treated with 1 mM H_2_O_2_ and number of viable bacteria was determined at the same time intervals. Bacterial colonies were counted after 48 h and expressed as log_10_ CFU/mL.

### Superoxide dismutase cloning and plasmid construction

After genomic DNA extraction by CTAB method (8), the *mlr7636* gene was amplified by polymerase chain reaction (PCR) using the specific forward primer SOD-XbaI-F (5’-ATAT<u>TCTAGA</u>CCACGAGGGAGTACTACCC**ATG**G-3’, the *XbaI* restriction site is underlined and the bold letters represent the start codon of *mlr7636* gene) and SOD-BamHI-R (5’-ATAT<u>GGATCC</u>TCACTTTGCCTTTTCGTAGAGC 3’, the *BamHI* restriction site is underlined), for directional cloning in pFAJ1708 downstream of the constitutive *nptII* promoter. PCR reaction was performed as follows: initial denaturation at 94°C for 3 min; 35 cycles at 94°C for 20 sec, annealing at 65°C for 20 sec and extension at 68°C for 1 min. The 639-bp PCR product and pFAJ1708 plasmid were digested by *XbaI* and *BamHI* enzymes, and DNA fragments were purified from agarose using Qiagen Gel Extraction QiaexII kit. The *mlr7636* gene was cloned into pFAJ1708 with T4 DNA ligase (Promega) at 4°C during 16 h. The resulting recombinant plasmid pFAJsod was transformed into *E. coli* DH5α using the heat shock method. Clones were sequenced to confirm the insertion of *mlr7636* in frame with *nptII* promoter, and used to transform Electrocompetent *E. coli* S17-1 by electroporation with a Gene Pulser Xcell, Bio Rad, 2.5 KV /5 ms.

### Transformation of *M. loti*

Biparental conjugation was performed in YEM medium using *M. loti* MAFF303099 (WT) as recipient and *E. coli* S17-1 carrying pFAJsod or pFAJ1708 as donors (9). The donor/recipient ratio was 1:1. Transconjugant *M. loti* cells were selected with Tc 20 μg/mL.

### Protein extraction

For the analysis of whole cell fraction, *M. loti* cultures were grown up to 0.1 OD_600_ in 50 mL liquid YEM medium. Cells were harvested by centrifugation at 15000 *g* at 4°C for 15 min, washed with 5 mL of 0.9% (W/V) NaCl and resuspended in 200 μl of 50 mM KH_2_PO_4_ pH 7.8 buffer. Cells were sonicated (Ultrasonic Vibracell VCX600) at 33% amplitude by pulses of 3 seconds with intervals of 3 seconds for 2 min. The samples were maintained in an ice-water bath during sonication. Extracts were centrifuged at 14000 g for 20 min and the supernatant was recovered. Periplasmic proteins were obtained by hypo-osmotic shock treatment (10). Cells were harvested by centrifugation at 15000 *g* and 4°C for 15 min, washed with 5 mL of 0.9% (W/V) NaCl and briefly resuspended in 0.2 mL of hyperosmotic buffer containing 20% (W/V) sucrose in 30 mM Tris/HCl pH 8.0 supplemented with 5 μl lysozyme (100 mg/mL in 30 mM Tris/HCl, pH 8). After 1 hour of incubation in ice, cells were pelleted by centrifugation for 5 min (10000 g at 4°C) and resuspended in 0.2 mL of ice-cold distilled water. After additional 15 min on ice, cells were pelleted by centrifugation at 10000 *g* at 4°C for 30 min. The supernatant contained the periplasmic proteins. Protein concentrations were determined by the Lowry method (11) using bovine serum albumin (BSA) as a standard.

### Polyacrylamide gel electrophoresis and western blotting

Sodium dodecyl sulphate polyacrylamide gel electrophoresis (SDS-PAGE) was performed as described by Laemmli (12). Non-denaturing polyacrylamide gel electrophoresis (ND-PAGE) was performed using the same buffer system with the omission of SDS from all buffers. Western blot analysis was conducted in a submerged system (Bio-Rad); the proteins separated by PAGE were blotted onto a nitrocellulose membrane (Amersham Biosciences). Polyclonal antibodies rabbit anti-FeSOD (Agrisera AS 06 125) and anti-MnSOD (N-20, Santa Cruz, SC18503) were used as primary antibodies. Goat anti-rabbit and rabbit anti-goat Alkaline phosphatase conjugates were used as secondary antibodies.

#### Enzyme activities

SOD activity was determined spectrophotometrically at 560 nm (13). One Unit of SOD was defined as the amount that inhibits the photoreduction of nitrotetrazolium blue chloride (NBT) by 50%. The reaction mixture was composed of 50 mM potassium phosphate pH 7.8 containing 777 μM methionine, 448 μM NBT, 0.54 μM EDTA and 3.32 μM riboflavin. In the absence of the enzyme, the mixture was calibrated to reach an OD_560_ = 0.25 after 10 min incubation in UV light (360 nm) at 25 °C. Specific activity was expressed in SOD units (μg of total protein)^−1^.

SOD zymogram was obtained after electrophoresis in 12% ND-PAGE. SOD activity was developed by incubating gels with 2.5 mM NBT during 25 min in dark, followed by incubation for 20 min in 50 mM potassium phosphate pH 7.8 containing 28 μM riboflavin and 28 mM tetramethyl-ethylene diamine (TEMED). The reaction was continued for 10 to 15 min under white light and then stopped by a brief wash in water (14).

To determine the metallic co-factor of SOD in zymogram, the ND-PAGE were pre-incubated with potassium cyanide (KCN 5 mM) or hydrogen peroxide (H_2_O_2_ 10 mM) at RT for 30 min in order to inhibit CuZnSOD and FeSOD, respectively. After incubation, the enzyme activity was assayed as described above.

Catalase (CAT) activity was determined with a reaction mixture containing 50 mM potassium phosphate pH 7.4 with the sample, and the reaction was started by adding 5 mM H_2_O_2_. Activity was determined spectrophotometrically by measuring the decreasing rate of H_2_O_2_ at 240 nm at 37°C for 1 min. Specific activity was expressed in CAT units (μg of total protein) ^−1^ (15).

SOD and CAT activities were evaluated in rhizobia grown in NaCl 150 mM and PEG 15%. Glucose 6 Phosphate Dehydrogenase (G6PDH) activity: Absence of cytoplasmic content in periplasmic fraction was evaluated by measuring G6PDH activity (16). The reaction mixture contained 100 mM Tris-ClH buffer pH 8, 10 mM MgCl_2_, 0.18 mM NADP^+^ and 1 mM D-Glucose-6-Phoshate. The reaction was initiated by adding the sample, and followed by formation of reduced NADPH at 340 nm in spectrophotometer at 25°C. One Unit of G6PDH was defined as the amount of enzyme required to catalyze the reduction of 1 μmole of NADP^+^ per min at 25°C. Specific activity was expressed in G6PDH units (mg of total protein)^−1^.

### Phylogenetic analysis

A phylogenetic tree was constructed with the Mlr7636 amino acid sequence based on the alignment of the 11 SOD sequences from the Rhizobase database sharing more than 50% of identity. The analysis was performed by the neighbor joining method (1000x bootstrap replications) using Mega 5.2 software (17).

### Bioinformatic analysis for signal peptide prediction

Protein sequence corresponding to the open reading frame of the locus *mlr7636* of *M. loti* MAFF303099 was obtained from genome.microbedb.jp/Rhizobase and was entered as query sequence in the SignalP 4.1 for Gram negative and TatP 1.0 algorithms (18, 19), using the default criteria.

### Statistical analysis

Values of enzymatic activity were expressed as mean ± standard error (SE). In all experiments, replicates were analyzed statistically by ANOVA using the DGC Test (Di Rienzo *et al*. 2002) at *p*≤ 0.05 in InfoStat software (20).

## Acknowledgements

This work was supported financially by Instituto Nacional de Tecnología Agropecuaria INTA, Argentina. Projects PNAGUA 1133032 and PNCYO 1127033.

